# Inclusion of maintenance energy improves the intracellular flux predictions of CHO

**DOI:** 10.1101/2020.12.21.423738

**Authors:** Diana Széliová, Jerneja Štor, Isabella Thiel, Marcus Weinguny, Michael Hanscho, Gabriele Lhota, Nicole Borth, Jürgen Zanghellini, David E Ruckerbauer, Isabel Rocha

## Abstract

Chinese hamster ovary (CHO) cells are the leading platform for the production of biopharmaceuticals with human-like glycosylation. The standard practice for cell line generation relies on trial and error approaches such as adaptive evolution and high-throughput screening, which typically take several months. Metabolic modeling could aid in designing better producer cell lines and thus shorten development times. The genome-scale metabolic model (GSMM) of CHO can accurately predict growth rates. However, in order to predict rational engineering strategies it also needs to accurately predict intracellular fluxes. In this work we evaluated the agreement between the fluxes predicted by pFBA using the CHO GSMM and a wide range of ^13^C metabolic flux data from literature. While glycolytic fluxes were predicted relatively well, the fluxes of tricarboxylic acid (TCA) cycle were vastly underestimated due to too low energy demand. Inclusion of computationally estimated maintenance energy significantly improved the overall accuracy of intracellular flux predictions. Maintenance energy was therefore determined experimentally by running continuous cultures at different growth rates and evaluating their respective energy consumption. The experimentally and computationally determined maintenance energy were in good agreement. Additionally, we compared alternative objective functions (minimization of uptake rates of seven nonessential metabolites) to the biomass objective. While the predictions of the uptake rates were quite inaccurate for most objectives, the predictions of the intracellular fluxes were comparable to the biomass objective function.

## Introduction

Chinese hamster ovary (CHO) cells are currently the leading production host for the synthesis of complex biopharmaceuticals with human-like post-translational modifications [1]. Products made in CHO belong to the top-selling drugs on the market (e.g. Humira^®^) [2]. The increasing demand for CHO-derived products requires advances in cell line and process development. Until now, significant improvements in productivity, product yield and growth rate of the cells have been achieved by media optimization and high-throughput screening for good producers [3,4]. However, the development of high producer cell lines is laborious, expensive and takes several months for each new product [5]. Systems biology approaches such as metabolic modeling might push the productivity even further, shorten the development times for new products and improve the product quality by elucidating potential bottlenecks in metabolism and suggesting genetic engineering or feed/media optimization strategies [6,7].

In 2016, a community-derived, consensus genome-scale metabolic model (GSMM) of CHO was published [8] and several updates have been made since [9–11]. These serve as a basis for applying genome-scale metabolic modeling to CHO. Simulations based on this model suggested huge potential for improved protein productivities [8]. Indeed, productivity was recently increased by implementing targeted knock-outs of several secreted host cell proteins. These results were consistent with the predictions of the CHO GSMM coupled with the secretory pathway model [11], which showed that these knock-outs would free up cellular resources [12].

To successfully use modeling for the rational design of new engineering strategies, accurate predictions of cellular phenotypes are essential. Previously we showed that a GSMM can accurately predict growth rates if supplied with accurate exchange rates of (essential) amino acids (AAs) and correctly determined biomass composition [13–15]. However, the model also needs to accurately predict intracellular fluxes. Previously it was shown that biomass composition [16] as well as extracellular exchange rates [17] have a big impact on the predicted intracellular fluxes. However, the validation of the flux predictions by the CHO GSMM with experimental data has been done only in one study so far [9].

In this work, we compare fluxes predicted by parsimonious flux balance analysis (pFBA), using the GSMM of CHO [8], against 20 ^13^C metabolic flux data sets across producer and non-producer cell lines in different media and culture modes (batch, fed-batch, semi-continuous) extracted from six publications [18–23]. We find that many fluxes in central carbon metabolism can only be reliably estimated if non-growth associated cellular maintenance is considered which was so far not included in the GSMM.

## Materials and methods

### Data and code availability

All data and scripts for the simulations and visualisation are available in Mendeley Data at https://data.mendeley.com/datasets/p973bk79ck/draft?a=12eceafb-76be-4ab5-8a53-86788fda6210.

### pFBA simulations

PFBA [24] was performed with the package COBRApy [25] using the solver Gurobi 9.1.0 [26] in python 3.7.9. The GSMM of CHO iCHO1766 [8] was used. Maximization of biomass production was used as the objective function (reactions `R_biomass_cho` and `R_biomass_cho_producing`). Uptake and secretion rates of extracellular metabolites and the recombinant product from 20 datasets (six publications) were used as constraints [18–23] (see Table S1 for an overview). In several cases, the uptake rates of tryptophan were not available. Assuming that it is the least abundant AA in the biomass [27], tryptophan uptake was constrained to the same value as the AA with the lowest uptake. All data were converted to mmolg^−1^ h^−1^ using dry cell masses provided in the publications. If not available, the average mass of CHO (264 pg) was used [13]. In some cases, the experimental data for the exchange rates was not provided, so the fitted values from ^13^C metabolic flux analysis (MFA) were used. If the data was provided only as plots, it was extracted using WebPlotDigitizer (https://apps.automeris.io/wpd/). Oxygen uptake was left unconstrained.

### Mapping of ^13^C models to *i*CHO1766

In order to compare predictions made by the GSMM of CHO to the results of ^13^ C MFA, it was necessary to map the metabolites and reactions from all models used for ^13^ C MFA (referred to as “^13^C models” in the further text) to the GSMM of CHO iCHO1766 [8]. Metabolic flux data was extracted from six publications [18–23], see Table S1 for an overview.

For most models, it was impossible to make a one to one mapping. Thus, the following rules were applied.

- If one reaction in a ^13^C model could be mapped to several reactions in *i*CHO1766, the fluxes from these reactions were summed up or subtracted, depending on the direction.
- In case of multiple equivalent reactions occurring in several compartments, their individual contributions predicted by pFBA were summed up and only the total was used for comparison. This approach disregards cellular compartments.
- In the ^13^ C models, several reactions are often lumped into one; therefore, the net flux of the corresponding reactions in the GSMM was calculated and compared to the flux of the lumped reaction.
- In case of producer cell lines, reactions for the synthesis of the recombinant protein were added comprising the AA composition provided in the publications and the energy demand for the polymerisation from [28] (2 GTP and 1.306 ATP per 1 mole of AAs are hydrolysed to 2 GDP, 1 AMP and 0.306 ADP).

### Computational estimation of the maintenance energy

To estimate maintenance energy (mATP), pFBA was run for every individual dataset, where growth rate was maximized and mATP hydrolysis (reaction `“R_DM_atp_”`) was constrained to a range of different values of mATP (0-40 mmol g^−1^ h^−1^ or until the simulation was no longer feasible). For each value of mATP, the agreement between experimental and predicted fluxes was evaluated and the value that lead to the lowest median relative error was chosen as optimal. Reactions with the experimental fluxes less than 1% of the maximum flux 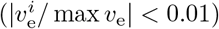 were omitted from this analysis, because their absolute fluxes were very small (often close to zero) and consequently the relative errors were very high (even though the absolute differences in fluxes were very small). This often distorted the analysis and no clear minimum was observed.

Additionally, the analysis was performed with all datasets at the same time – the mATP value was varied and the overall agreement between the predicted and the experimental fluxes for all datasets was evaluated. The calculated median errors were divided by the number of datasets for which a feasible solution was obtained (because the higher the mATP value, the lower the amount of datasets with a feasible pFBA solution).

### Uptake objective function

PFBA simulations were done as in [29]. First, growth rate and productivity were fixed to the experimental values and the uptake of each essential AA was minimized one by one to get an estimate for the minimum uptake rate that sustains the experimental growth rate (nine AAs are essential in the iCHO1766 model). If the measured experimental uptake rate was lower than the minimum required uptake by the model, we fixed it to the computed uptake. Otherwise the experimental uptake rate was used. In a few cases, the uptake rates of tyrosine and cysteine had to be adjusted in the same way as the essential AAs, because the experimental uptake rates were insufficient to sustain growth and no solution was obtained (three datasets for tyrosine and three for cysteine).

In the next step, we constrained the nonessential uptake rates, secretion rates, growth rate and productivity to the experimental values (except for the nonessential uptake rate that was set as the objective, which was left unconstrained) and the essential uptake rates to the predicted or experimental values (see above). The flux distributions were obtained by performing pFBA with minimization of an uptake rate (glucose, glutamine, serine, tyrosine, arginine, aspartate or asparagine) as the objective.

### Statistical analysis

Statistical analysis was carried out in R version 4.0.2. Linear correlations between experimental and predicted data were calculated with R function `lm`. In case of intracellular flux predictions, the inverses of the experimental confidence intervals were used as weights for the linear fitting. The relative error of the fluxes was calculated with Eq (1),

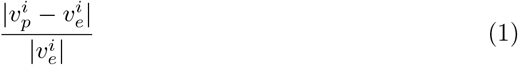

where 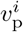 are the fluxes predicted by pFBA and 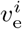 are the experimentally determined fluxes. Mean and median relative error were calculated.

### CHO cell cultivation

Suspension CHO-K1 cells (ECACC CCL-61) were grown in CD-CHO medium (Gibco™ Thermo Fisher Scientific, MA, USA) supplemented with 0.2% (v/v) Anti-Clumping Agent (Gibco™ Thermo Fisher Scientific, MA, USA) and 8 mM L-Glutamine (Sigma-Aldrich, MO, USA). Cells were cultivated in 125 mL non-baffled Erlenmeyer flasks at 37 °C at 140 rpm with 25 mm throw, 7% CO_2_ and 85% humidity and passaged every 2-4 days. Mycoplasma contamination was regularly checked with MycoAlert™ Mycoplasma Detection Kit (Lonza, Basel, Switzerland). The cell concentration and viability were determined with Vi-CELL XR (Beckman Coulter, CA, USA) calibrated with ViaCheck™ Concentration Control (Bangs Laboratories, Inc., IN, USA).

### Continuous cultivation

Cells were cultivated in DASGIP^®^ Parallel Bioreactor System (Eppendorf, Hamburg, Germany) in DS0700ODSS vessels at 37 °C and agitated with a marine impeller at 80 rpm. pH was monitored with EasyFerm Plus PHI K8 225 pH Electrode (Hamilton, NV, USA) and maintained at 7 ± 0.05 with CO_2_ and 7.5% (w/w) NaHCO_3_. pH was also checked with an external pH probe (Mettler Toledo) at least twice per week to correct potential pH drifts. Dissolved oxygen was measured with DO sensor VisiFerm (Hamilton, NV, USA) and maintained at 30% with a cascade (1. increase O_2_ concentration in the incoming gas up to 50%, 2. increase both flow rate and O2 concentration up to 75%, 3. increase flow rate up to 0.1 volume per volume per minute [vvm]). Cells were inoculated at a seeding density of 1.6 × 10^5^ viable cells/mL at a working volume of 300 mL. At the end of the exponential phase, the culture was switched to the continuous mode and maintained at a constant volume of 270 mL. The flow-in pump was set to a constant rate and the amount of medium pumped into the bioreactors was monitored using Mettler Toledo balances MS6002TS (readability 0.01 g) to calculate an accurate flow rate into the bioreactors. The tube for flow-out was positioned at the height corresponding to 270 mL and the flow rate was set to a higher value than flow-in to prevent overflow. The feed medium was kept at room temperature and protected from light with aluminum foil. Due to the instability of glutamine, the medium was exchanged every 5-7 days. During this time frame, the glutamine degradation was shown to be negligible (see Fig S6).

After changing the dilution rate, the cultures were left to equilibrate for at least five volume exchanges (except for one dilution rate (DR3, see Table 1) which was interrupted because of contamination). To verify whether the cultures reached steady state, linear fits were performed with viable cell density and Bioprofile data (glucose, lactate and ammonium concentrations) using R function `lm`. If the 95% confidence interval of the slope contained zero, the parameter was considered stable. For seven out of eleven dilution rates, all parameters were stable, including a dilution rate where only 2.5 volume exchanges were reached due to contamination (DR3). For three dilution rates 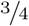 parameters were stable, but the concentration changes of the unstable parameters were within the measurement error of the measurement device. Overall, all dilution rates were deemed stable enough and were used for further analysis (see Table 1 for an overview).

**Table 1.** The dilution rates, calculated growth rates (Eq (3)) and the steady state concentrations of cells and metabolites.

| ID | Dilution rate [h<sup>-1</sup>] | Growth rate [h<sup>-1</sup>] | Viable cells mL<sup>-1</sup> 10<sup>-6</sup> | Glucose [mmol g<sup>-1</sup>h<sup>-1</sup>] | Lactate [mmol g<sup>-1</sup>h<sup>-1</sup>] | Ammonium [mmol g<sup>-1</sup>h<sup>-1</sup>] |
| --- | --- | --- | --- | --- | --- | --- |
| DR1 | 0.020 | 0.021 | 5.43 | -0.42* | 0.56 | 0.08 |
| DR2 | 0.026 | 0.027 | 5.42 | -0.44 | 0.53 | 0.09* |
| DR3 | 0.016 | 0.016 | 6.02 | -0.32 | 0.31 | 0.06 |
| DR4 | 0.027 | 0.028 | 6.06 | -0.43 | 0.58 | 0.09 |
| DR5 | 0.032 | 0.032 | 6.31 | -0.48 | 0.57 | 0.11 |
| DR6 | 0.033 | 0.033 | 5.83 | -0.46 | 0.61 | 0.12 |
| DR7 | 0.032 | 0.033 | 5.32 | -0.46 | 0.70 | 0.12 |
| DR8 | 0.023 | 0.023 | 11.37 | -0.29 | 0.26 | 0.04 |
| DR9 | 0.035 | 0.036 | 6.14 | -0.54** | 0.62** | 0.09** |
| DR10 | 0.020 | 0.020 | 10.24 | -0.28 | 0.27 | 0.04 |
| DR11 | 0.024 | 0.024 | 9.77 | -0.35* | 0.34 | 0.04 |

A parameter was considered stable when the 95% confidence interval of the slope from the linear fit contained zero.
\* The slope was statistically significant, but the change in concentration of glucose and ammonium was within the measurement error of the Bioprofile analyzer.
\*\* Confidence intervals could not be calculated because only two data points were available. The change in concentration was within the measurement error of the Bioprofile analyzer.

### Extracellular metabolites

The samples for supernatant analysis were taken at least at two time points per steady state. To separate cells from the medium, the cultures were centrifuged for 8 min at 200 g at room temperature and supernatants were stored at −80 °C until further analysis or processed immediately. Lactate, ammonium and glucose concentrations were measured at Bioprofile 100Plus (NOVA Biomedical, MA, USA). Lactate and ammonium measurements were corrected with 4-point calibration curves made in CD-CHO medium. As glucose was already contained in the CD-CHO medium, a calibration curve was not done. Instead, the average measured concentration of the standards was used for the calculation of the uptake rates in Eq (4).

For two dilution rates (DR1 and DR2), the AA concentrations were quantified by Biocrates with a commercial AbsoluteIDQ^®^ p180 kit. Briefly, AAs were derivatized with phenyl isothiocyanate in the presence of internal standards. The quantification was performed by liquid chromatography-mass spectrometry (LC-MS/MS) using a 4000 QTRAP^®^ (AB Sciex, Darmstadt, Germany) and a Xevo TQ-S micro (Waters, Vienna, Austria) instrument with an electrospray ionization source.

For the remaining dilution rates, AAs were analyzed using a high-performance liquid chromatography method. Briefly, samples were diluted, internal standards 3-(2-thienyl)-DL-alanine (Fluka) and sarcosine (Sigma) were added and subsequently filtered using a 0.2 *μm* filter unit (Sartorius). In an automated pre-column derivatization method, free primary AAs reacted with ortho-phthalaldehyde (OPA, Agilent) and proline and hydroxyproline with 9-fluorenyl-methyl chloroformate (FMOC, Fluka) and were then separated on a ZORBAX™ Eclipse Plus C18 column (Agilent) at 40 °C using a flow rate of 0.64 mL/min. After gradient elution with 10 mM K_2_HPO_4_:10mM K_2_B_4_O_7_ (Merck) pH 8.2 as solvent A and acetonitrile:methanol:water (45:45:10, v:v:v) (Merck) as solvent B, AAs were excited at 230 nm and the fluorescence signal was detected at 450 nm for OPA derivates and 266 nm and 305 nm for FMOC derivates, respectively. Samples were quantified using an internal standard calibration.

The metabolite concentrations in the medium measured by both methods were compared to the patent for CD-CHO medium (Gibco™ Thermo Fisher Scientific, MA, USA) to make sure the results are in the expected range and comparable between the two methods.

The bioprocess and metabolite concentration data is available in Mendeley Data at https://data.mendeley.com/datasets/p973bk79ck/draft?a=12eceafb-76be-4ab5-8a53-86788fda6210.

### Calculation of growth rate and exchange rates

For all calculations, the average of all data points in each steady state was used. Dilution rate *D* was calculated with Eq (2),

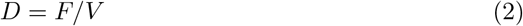

where *F* is the flow rate into the bioreactors (calculated from the change of mass of the fresh culture medium over time) and *V* = 270 mL is the volume of the medium in the bioreactors. The growth rate *μ* at steady state was calculated with Eq (3),

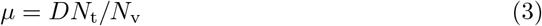

where *N*_v_/*N*_t_ denotes fraction of viable cells. Steady state exchange rates *q^i^* of extracellular metabolites were calculated with Eq (4),

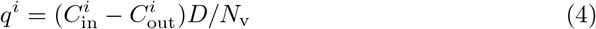

where 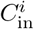 and 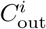 are concentrations of metabolites in the incoming medium and in the bioreactor, respectively. To calculate standard deviation (SD) for an exchange rate, the SDs were calculated for each variable from all available data points in steady state. Then, the SD (*σ*) of the rate was calculated according to the mathematical rules of manipulation with standard deviations - Eq (5) if values were multiplied or divided (e.g. C = AB),

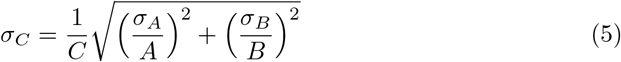

and Eq (6) if they were summed or subtracted (e.g C = A+B)

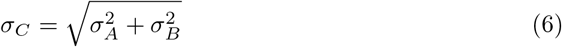

### Determination of maintenance energy

The GSMM with cell line specific biomass composition from [13] was used (iCHO_K1par-8mMCD, BioModels ID: MODEL1907260016). The experimentally determined growth rate and the exchange rates of glucose, lactate, ammonium and AAs were used as constraints for flux balance analysis (FBA). The lower and upper bounds of the exchange reactions and the biomass reaction (`“R_biomass_specific”`) were fixed to the experimental values ± SD. Due to high experimental noise, the uptake rates of some essential AAs were too low to sustain the experimental growth rate. Therefore the minimal uptake requirements were estimated as in [29] and, if necessary, the constraints were adjusted. This had no impact on the calculations of the ATP consumption as these AAs are solely used for biomass generation and not for ATP generation. In three cases, the lower bounds of secretion rates were relaxed (DR1 - alanine secretion by 25%, DR5 glycine secretion by 40%, DR6 aspartate by 25%), otherwise the solutions would have been infeasible (the upper bounds were set to the experimental values).

ATP hydrolysis was maximized (reaction `“R_DM_atp_”`) and the maximum ATP consumption was plotted against the growth rate. A linear fit was done in R (function `lm`) and the intercept represented the non-growth associated energy consumption.

## Results

### pFBA underestimates intracellular fluxes in *i*CHO1766

20 ^13^C MFA datasets from producer and non-producer CHO cell lines across different media were collected from literature [18–23], see Table S1. Flux data were mapped onto the genome-scale metabolic model iCHO1766 [8] and compared to pFBA predictions based on biomass maximization (see methods).

For 6 out of the 20 datasets growth could be predicted with an error of less than ±25% (for `R_biomass_cho`), see Fig 1a. In three datasets growth was vastly underestimated, and overestimated in the remaining eleven. In one extreme case growth was overestimated by 190%. Overall the median relative error was close to 60%. pFBA performs even worse when compared to measured intracellular fluxes, hitting an overall median relative error of 83.8% (Fig 1b). On average, fluxes in glycolysis and AA are predicted better than fluxes in pentose phosphate pathway (PPP), pyruvate metabolisms, and tricarboxylic acid cycle (TCA). More specifically, fluxes in PPP and TCA were vastly underestimated (median error 86.9% and 94.5%, see Table 2).

**Fig 1.**
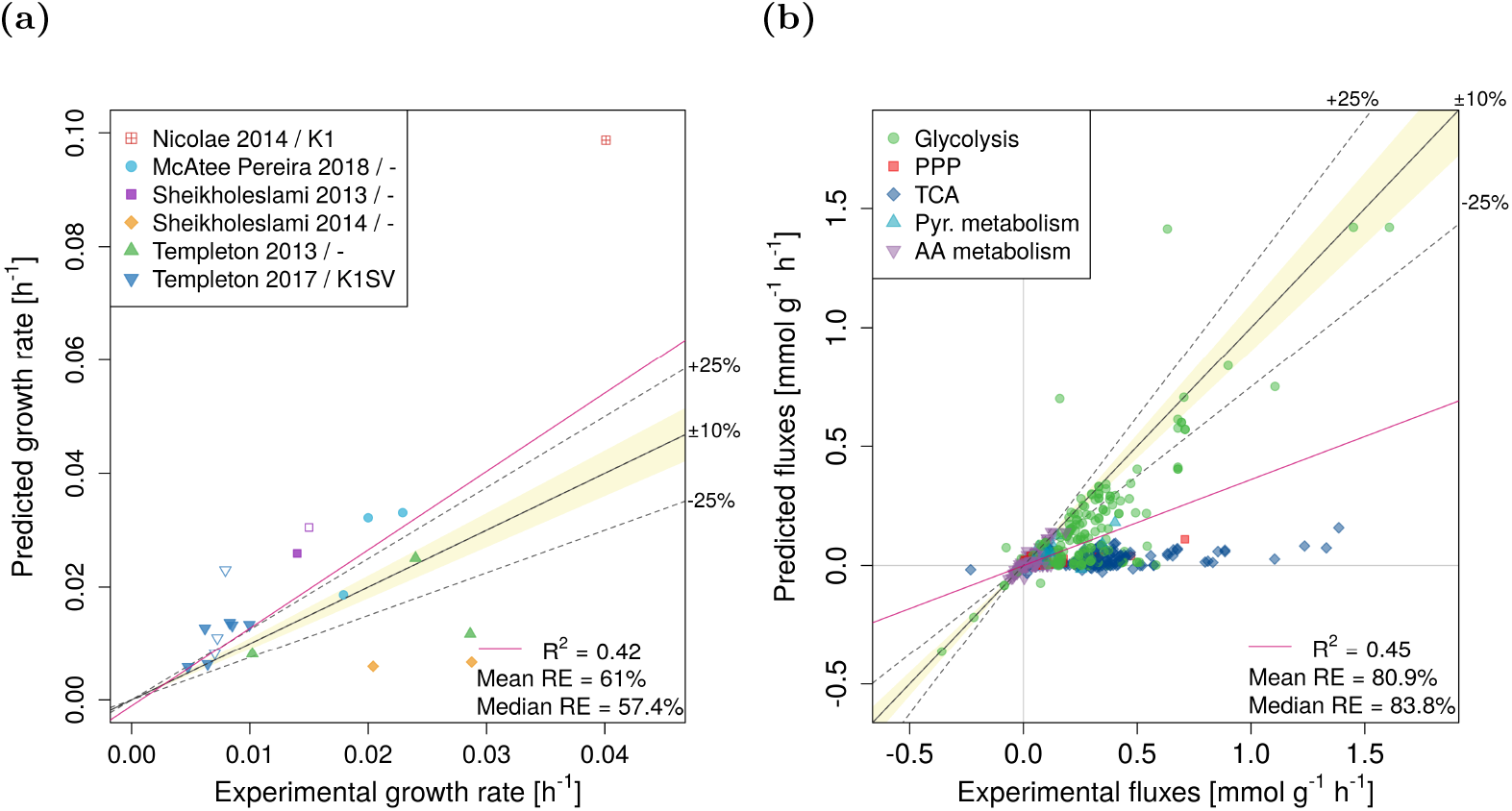
Experimental vs. predicted growth rates (panel a) and intracellular fluxes (panel b). Data is shown for biomass equation `R_biomass_cho` as the objective function. RE – relative error. The legend in panel a indicates the publication and the used CHO cell line (if the information was available). Empty symbols indicate non-producers.

**Table 2.**
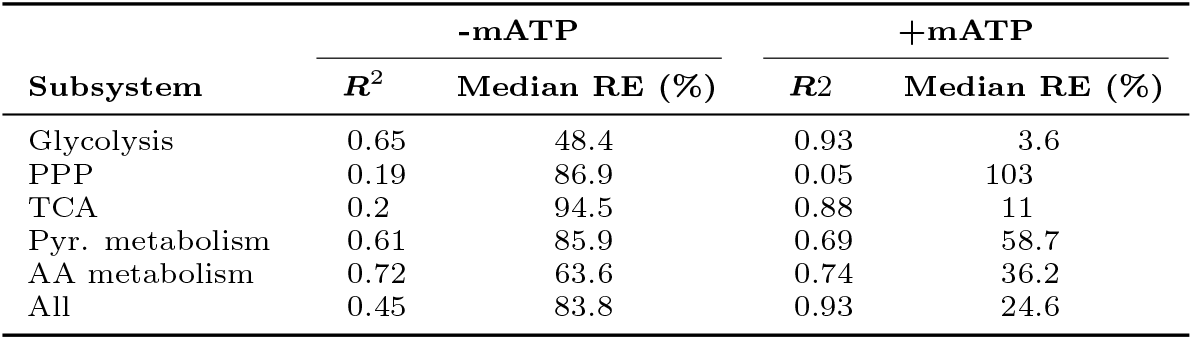
R^2^ and median relative error (Median RE) of the experimental and predicted fluxes with (+mATP) or without mATP (-mATP) as constraint. Data is shown for biomass equation `R_biomass_cho`.

| Subsystem | -mATP |  | +mATP |  |
| --- | --- | --- | --- | --- |
| | $R^2$ | Median RE (%) | $R^2$ | Median RE (%) |
| Glycolysis | 0.65 | 48.4 | 0.93 | 3.6 |
| PPP | 0.19 | 86.9 | 0.05 | 103 |
| TCA | 0.2 | 94.5 | 0.88 | 11 |
| Pyr. metabolism | 0.61 | 85.9 | 0.69 | 58.7 |
| AA metabolism | 0.72 | 63.6 | 0.74 | 36.2 |
| All | 0.45 | 83.8 | 0.93 | 24.6 |

We checked whether flux predictions can be improved when experimental growth rates were used as additional constraints. Yet, no improvement was observed for those datasets that returned a feasible solution (median relative error 91.5% vs. 83.8% previously).

### Maintenance energy improves predicted glycolytic and TCA fluxes

The gross underestimation of intracellular fluxes, especially in glycolysis and TCA (supplementary Figure S3), which are the major sources of ATP, points at an underestimation of the actual energy demand. In fact, current CHO models typically lack non-growth associated maintenance energy demands conventionally used in microbial models [30–32]. Thus, for each dataset we determined a CHO-specific, non-growth associated mATP by fixing the flux for the maintenance reaction `R_DM_atp_c_`) such that the median relative error across all fluxes was minimal. Fig S2 illustrates data for the cell line SV-M3 [22]. For this cell line we find a maintenance demand of 5.75mmolg^−1^ h^−1^. Across all cell lines, mATP averages at 5.9 and 6.4mmolg^−1^ h^−1^ for `R_biomass_cho` and `R_biomass_cho_producing`, respectively (Fig 2a). Again the non-producing cell line CHO-K1 [20] sticks out with a more than three times larger maintenance energy demand compared to the average (across cell lines) for `R_biomass_cho_producing`.

**Fig 2.**
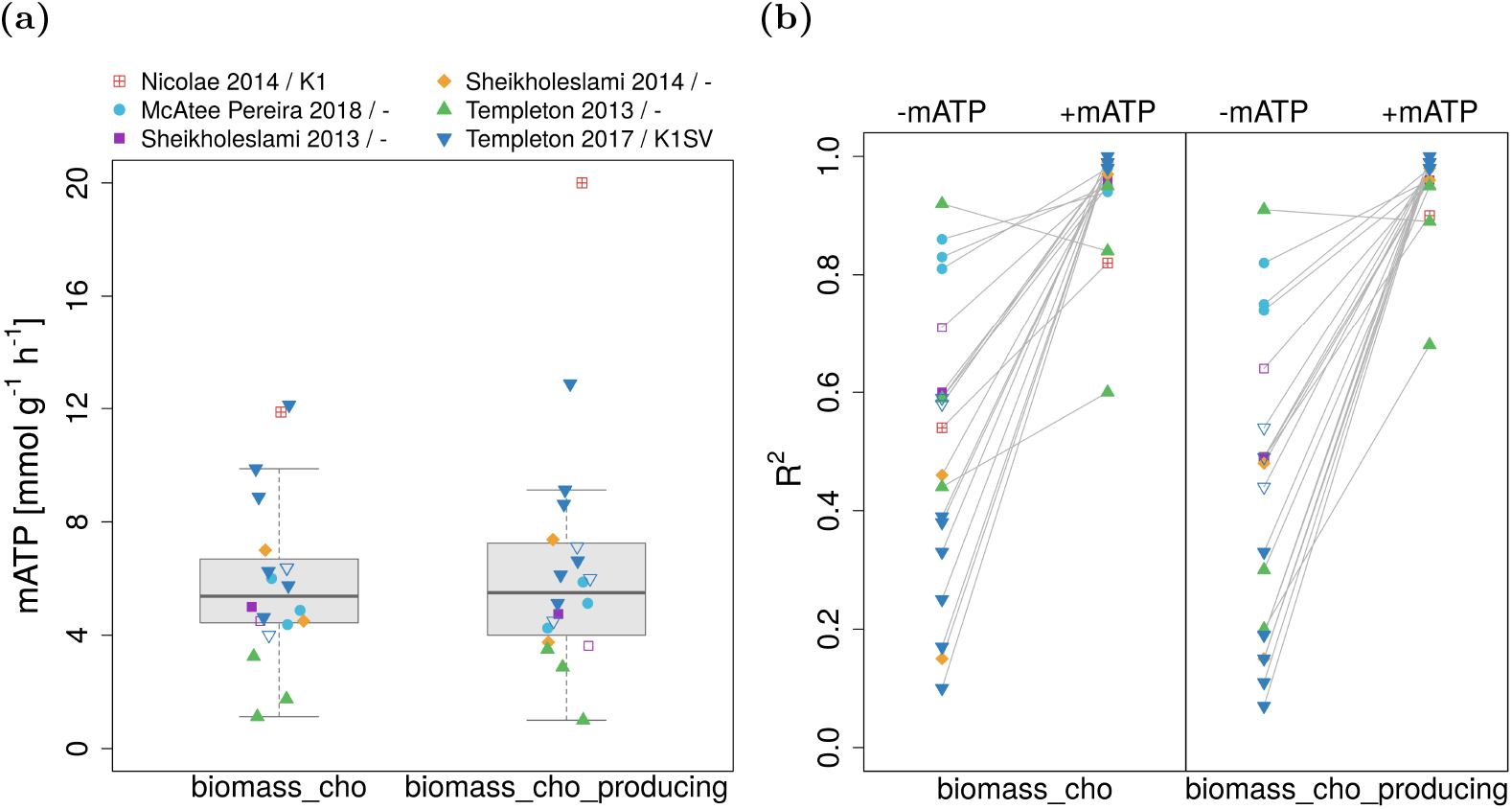
Estimated mATP values and their effect on flux prediction accuracy. Panel a: Computationally estimated mATP values for different datasets with two biomass equations. Panel b: R^2^ values from linear fits of experimental and predicted intracellular fluxes without (-mATP) or with mATP (+mATP) as constraint. The legend indicates the publication and the used CHO cell line (if the information was available). Empty symbols indicate non-producers

Additionally we estimated a mean maintenance energy by fitting mATP across all datasets. Mean maintenance was very close to the average mATP determined for each individual dataset (5.75 for both biomass equations vs. 5.9 and 6.4mmolg^−1^ h^−1^ for `R_biomass_cho` and `R_biomass_cho_producing`, respectively).

Enforcing a minimal mATP strongly decreases the prediction errors in the intracellular fluxes for all but one dataset, see Fig 2b. More specifically, the overall median relative error decreased from 83.8% to 24.6% (Fig 3b) and from 92.5% to 16.6% for `R_biomass_cho` and `R_biomass_cho_producing`, respectively. Conversely, *R^2^* more than doubled from 0.45 and 0.41 to 0.93 and 0.95.

**Fig 3.**
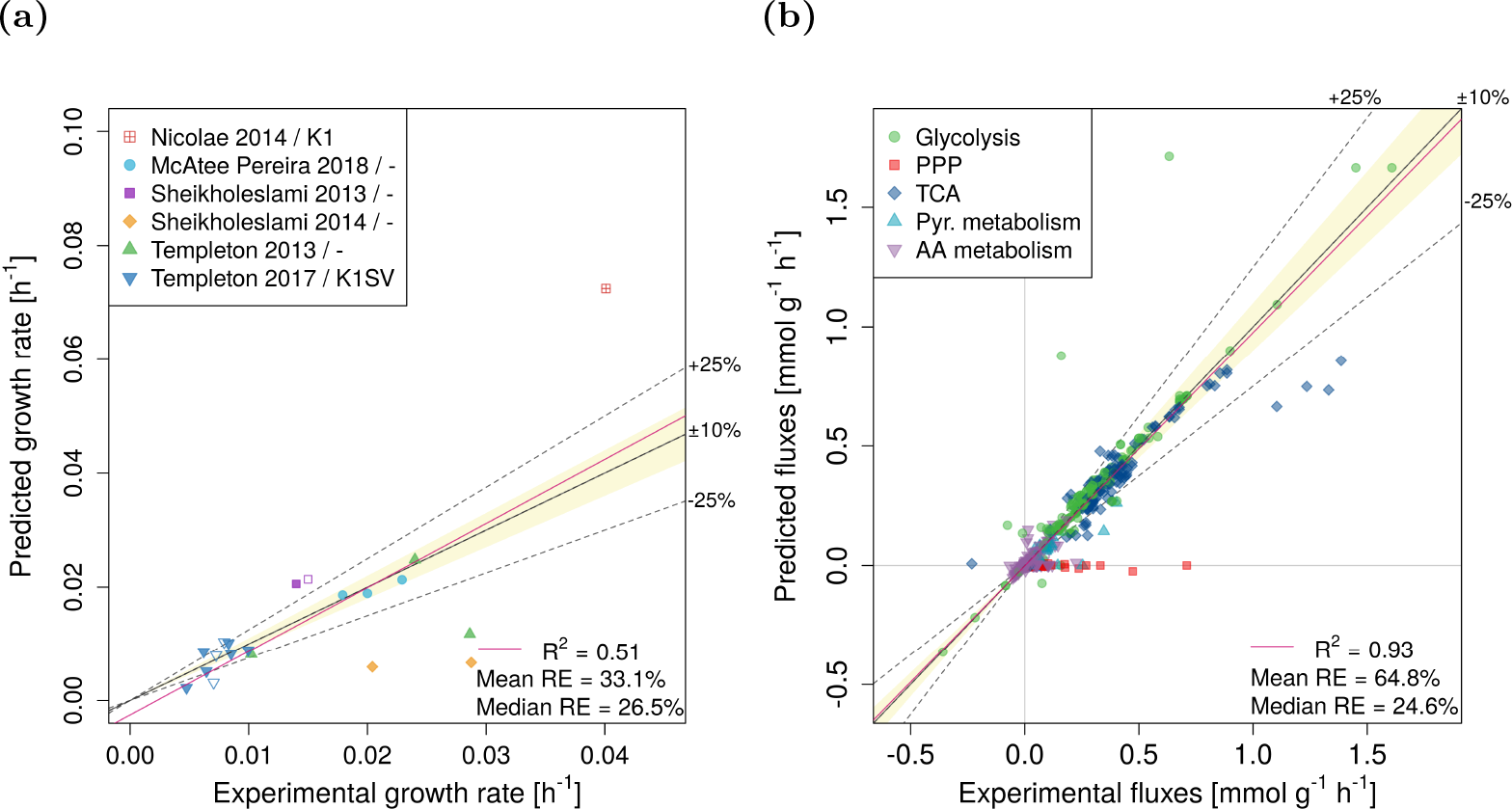
Experimental vs. predicted growth rates (panel a) and intracellular fluxes (panel b) after the addition of mATP as constraint. Results are shown for `R_biomass_cho` as the objective function. RE – relative error. The legend in panel a indicates the publication and the used CHO cell line (if the information was available). Empty symbols indicate non-producers

Not only intracellular fluxes, but also growth rates were better predicted (Fig 3a). Now ten rather than previously only six out of 20 growth rates could be predicted with an error of less than ±25%. Five were even predicted within an error band of ±10%.

While the predictions largely improved for TCA and glycolysis fluxes, PPP became inactive and the agreement with experimental data became worse (Table 2). However, the experimental data for PPP often have very big uncertainty – e.g. for Templeton 2013 [21], Templeton 2017 [22] and McAtee Pereira 2018 [23], which make 15 out of 20 datasets, the confidence intervals for the PPP reactions include zero.

### Minimizing non-essential nutrient uptake performs similar to maximizing growth

Recently, minimizing uptake of non-essential nutrients (rather than maximizing growth) was suggested to be a more suitable modeling objective for CHO [29]. Thus, we repeated all previous simulations with uptake objective function (UOF), using the exchanges of glucose, glutamine, serine, tyrosine, asparagine, aspartate and arginine as objectives. Maintenance energy was estimated as before and similar mean mATP values were obtained (5.6-6.4 and 5.8-6.7 for the biomass equations `R_biomass_cho` and `R_biomass_cho_producing`, respectively).

The prediction accuracy of the intracellular fluxes after the addition of the mATP constraint was comparable with biomass objective function (BOF) for all objective functions (see Fig 4b for an example with glucose UOF and Fig S4 for the remaining objectives).

**Fig 4.**
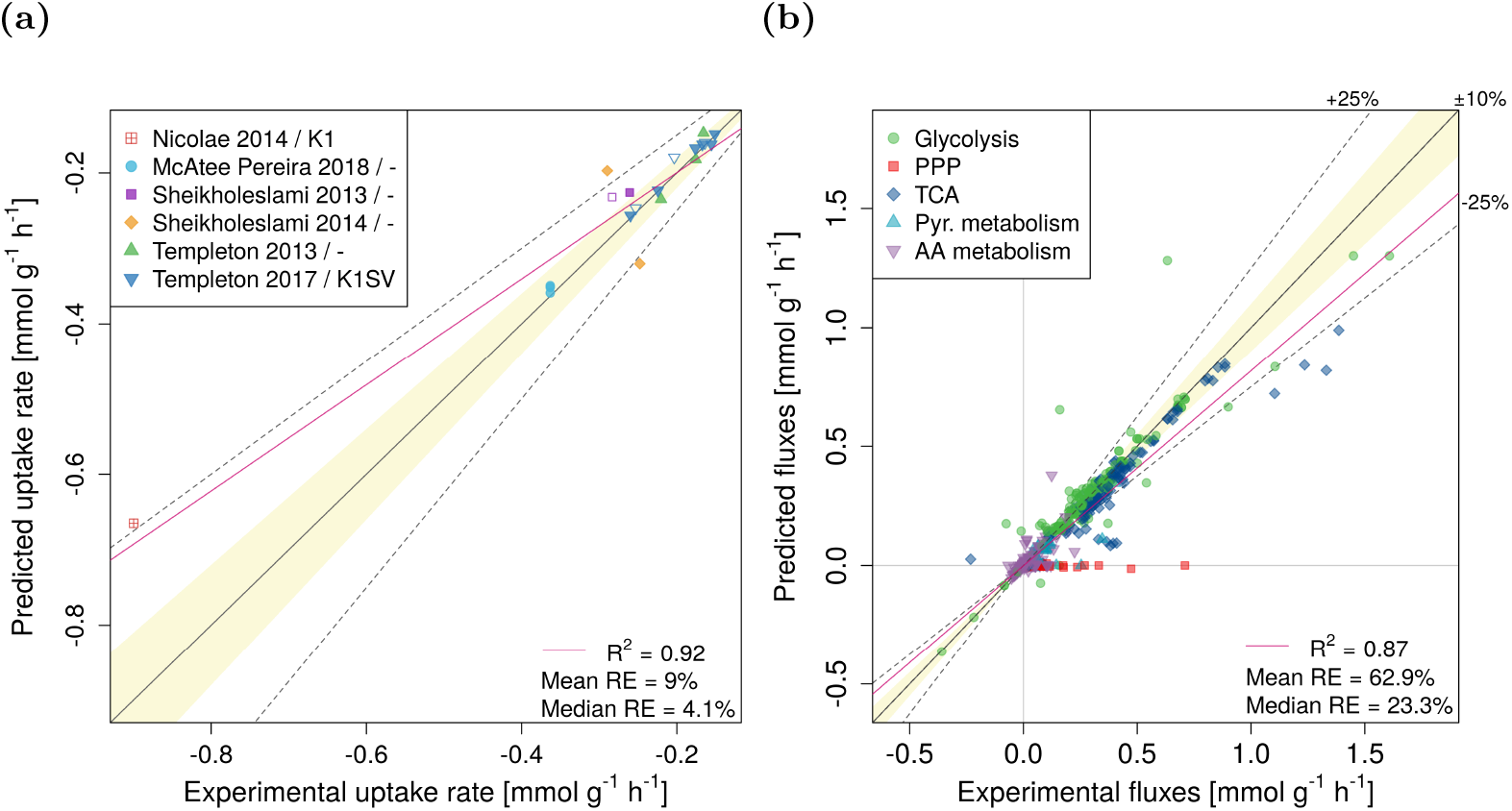
Experimental vs. predicted fluxes using minimization of glucose uptake rate as the objective function. Panel a: the experimental vs. predicted minimal glucose uptake rate. Panel b: experimental vs. predicted intracellular fluxes. Results are shown for `R_biomass_cho` as the biomass reaction. RE – relative error. The legend in panel a indicates the publication and the used CHO cell line (if the information was available). Empty symbols indicate non-producers

The predictions of the minimum uptake rates were best for glucose with *R*^2^ =0.92 and a median relative error of 4.1% (Fig 4a), followed by glutamine (*R*^2^ = 0.75, median error 50.4%) and asparagine (*R*^2^ = 0.12, median error 24.2%). However, the uptake rates of the remaining AAs were not predicted well (*R*^2^ = 0.06 or less; median errors within a range of 65.8-176%; Fig S4).

### Experimental determination of maintenance energy

To verify our computational estimate, we determined the maintenance energy experimentally in a CHO-K1 cell line. Continuous cultivation was run at eleven different dilution rates ranging from 0.016 to 0.035 h^−1^ (Table 1). Cell viability was above 95% for all steady states. The steady state viable cell densities were between 5.3 and 6.4 × 10^6^ viable cells/mL for eight dilution rates; for the remaining three they reached between 9.7-11.4 × 10^6^ viable cells/mL (Fig S7). For each dilution rate, extracellular exchange rates of glucose, AAs, lactate and ammonium were determined. Uptake of glucose, and glutamine as well as secretion of lactate and ammonium increased with increasing growth rate, see Fig 5. However, in case of waste product secretion rates, the three dilution rates that had higher steady state cell concentrations seem to separate from the remaining ones (indicated as magenta triangles in Fig 5).

**Fig 5.**
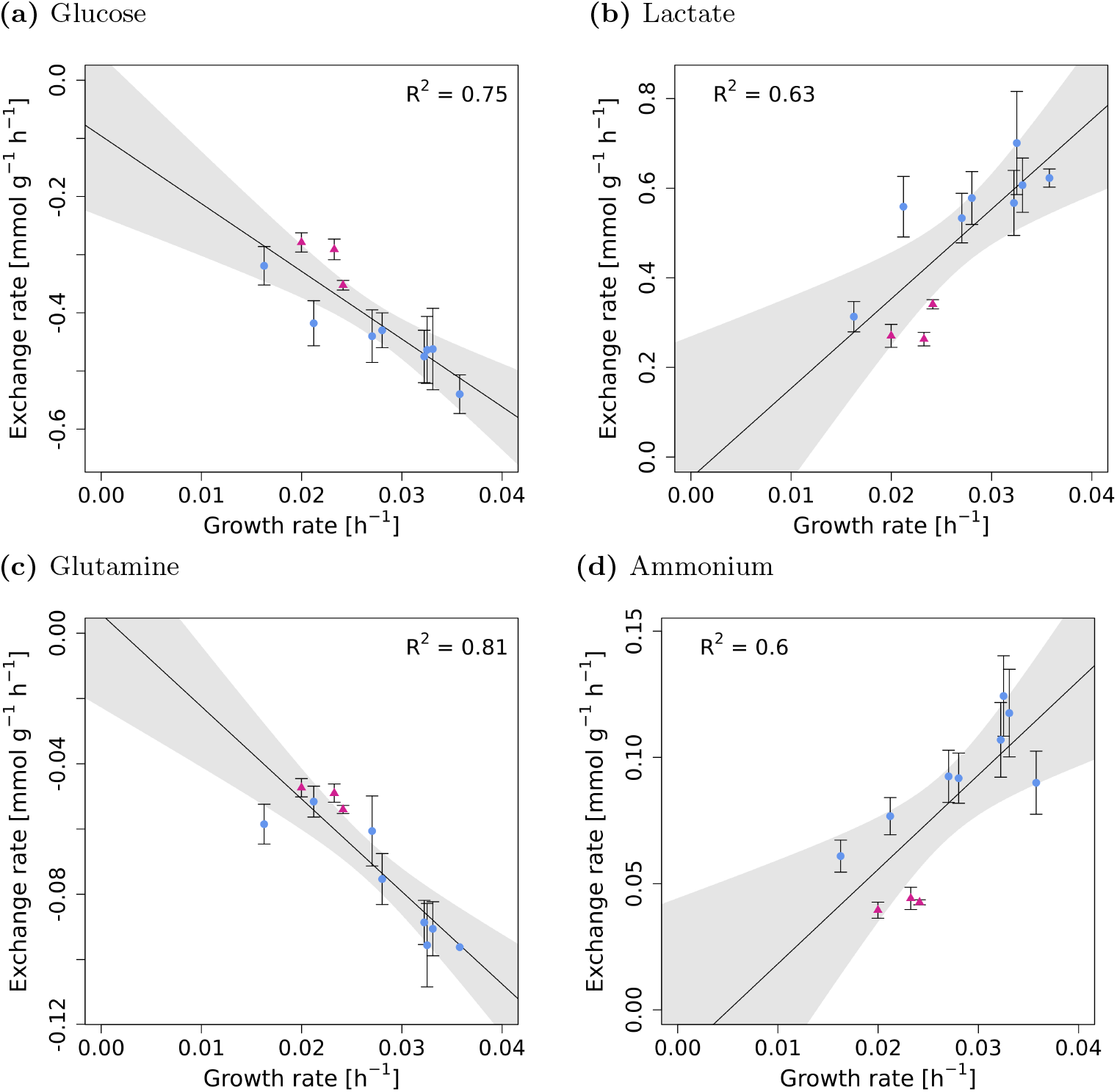
The experimental exchange rates of glucose (panel a), lactate (panel b), glutamine (panel c) and ammonium (panel d) increase with increasing growth rate. The shaded areas represent 95% confidence intervals. The triangle points in magenta color are the dilution rates that had unusually high cell concentration in steady state (see Fig S7).

Exchange rates of glucose, lactate, ammonium, all AA, and the growth rate were used as constraints for FBA and the hydrolysis of ATP was maximized. The ATP consumption was plotted against growth rate and a linear model was fitted (Fig 6). The intercept represents the non-growth associated ATP consumption - the estimated mATP and its standard error was determined to be 4 ± 1.6 mmol g^−1^ h^−1^, which compares well with the average mATP of 5.9/6.4 mmol g^−1^ h^−1^ determined computationally above.

**Fig 6.**
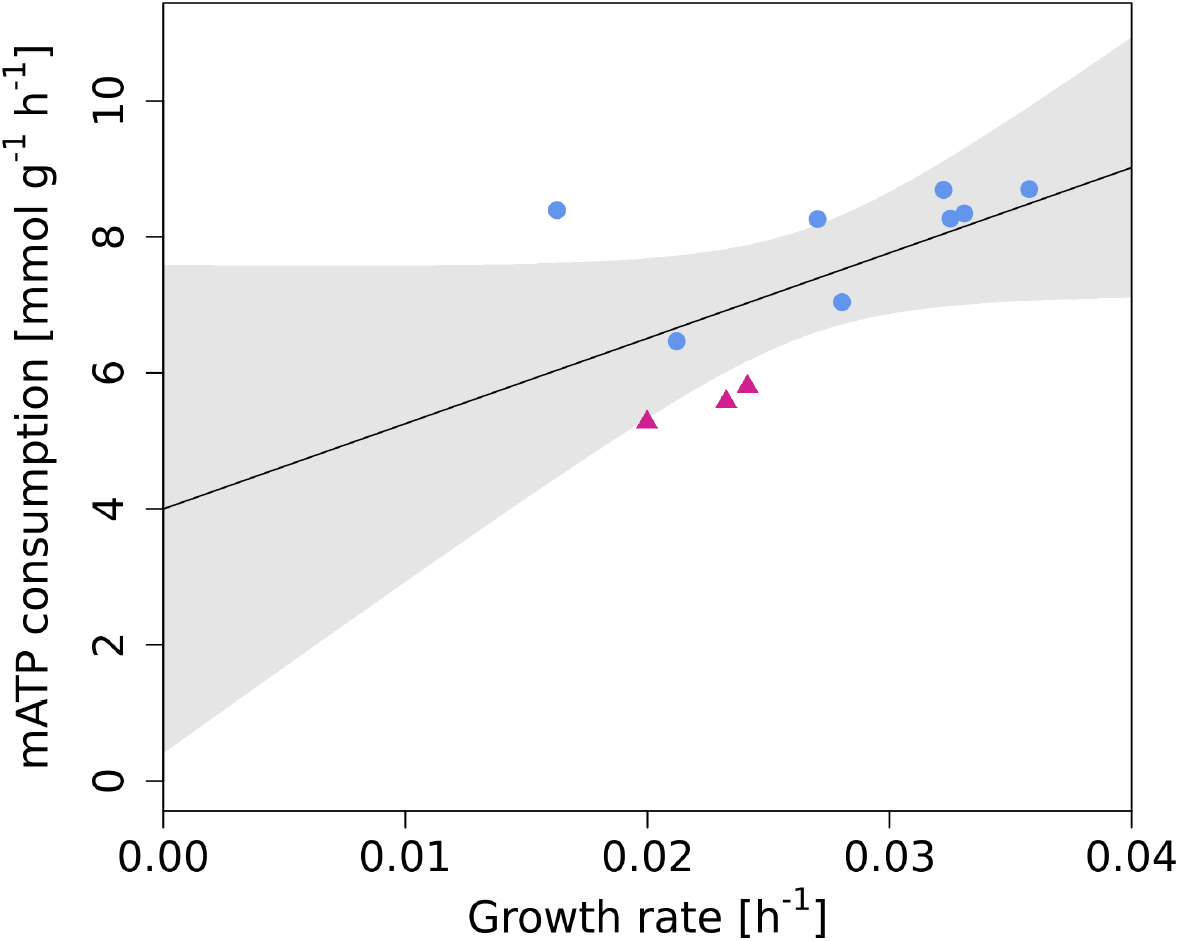
The rate of ATP consumption at different growth rates. The black line is a linear fit and the intercept represents the ATP consumption at zero growth rate. The magenta triangles are dilution rates that had unusually high cell concentration in steady state (see Fig S7). The shaded area represents 95% confidence interval.

## Discussion

An accurate determination of intracellular fluxes is key for understanding cellular metabolism and applying methods that predict engineering strategies. Intracellular fluxes can be experimentally determined with ^13^C metabolic flux analysis [33]. However, this method is very expensive due to the usage of labelled substrates and prone to experimental variability because of the need for rapid sampling and quenching of the metabolism. One of the cheaper and simpler methods for flux determination is parsimonious flux balance analysis (pFBA) [24], which first maximizes the biomass production and subsequently minimizes the total sum of fluxes, based on the assumption that cells try to minimize the utilization of resources. This was shown to be consistent with experimental data and it can be applied to genome-scale metabolic models, which can provide a more complete picture about cell metabolism than the small models used for ^13^C MFA.

In this work, we evaluated the agreement between experimentally measured intracellular fluxes from 20 datasets [18–23] and pFBA predictions made with iCHO1766 genome-scale model. We observed that the fluxes of central carbon metabolism, especially TCA cycle, were underestimated in all datasets, which was explained by an insufficiently represented energy demand in the model. Although the iCHO1766 model takes into account energy demands for the synthesis of biomass and recombinant proteins, it currently lacks a value for non-growth associated maintenance energy – the energy needed for processes such as turnover and repair of macromolecules or maintenance of concentration gradients (e.g. Na^+^ /K^+^ and Ca^2+^ ATPases) [34]. As no such value was available for CHO until now, we determined mATP computationally and experimentally.

The variability of the computationally estimated mATP across cell lines and conditions was quite high (relative SD 49% and 64% for `R_biomass_cho` and `R_biomass_cho_producing`, respectively). This might be the result of the experimental errors of the metabolite exchange rates. As seen in Fig 1a, the growth rate predictions had a high error, which we have shown previously to be sensitive to errors in the exchange rates [13,14]. Another factor is the error of the ^13^C flux measurements which often had considerably big confidence intervals. However, the differences might also stem from differences in the cell lines, cultivation conditions or productivities. Nevertheless, the average mATP values were very similar when estimated with different biomass equations (Fig 2a) and lead to a major improvement in the predicted intracellular fluxes, especially in the TCA and glycolysis. Fluxes of PPP got worse after the addition of a mATP constraint, which points at alternative NADPH sources connected to the TCA, e.g. malic enzyme. Indeed we found higher activity of malic enzyme in some datasets, but not consistently in all. This points to a possible lack of actual NADPH demand in the model – e.g. for protein folding or degradation of misfolded proteins [35].

We also investigated the effect of alternative objective functions (nonessential uptake rates) suggested by Chen et al. [29]. The estimated mATP values and the predictions of intracellular fluxes were comparable to the predictions done with the ‘‘traditional” BOF for all tested objectives. However, the predictions of the minimum uptake rates worked much better for glucose uptake rate compared to the AA uptake rates.

The choice of the appropriate objective function might depend on the availability of experimental data. In case of using BOF, highly accurate uptake and secretion rates are needed in order to obtain accurate predictions, especially for essential AAs [13,14]. If these are not available, using the UOF (glucose) might be a better choice than the use of BOF with imprecise AA uptake rates as constraints.

The experimentally determined mATP was comparable to the computational estimate, although the uncertainty was quite high due to the technical difficulty of running continuous fermentation and the unstable nature of CHO cells. Long cultivations lead to cell clumping, which complicated cell counting. It is also known that CHO cells are unstable during long term cultivations [36,37]. Furthermore, the physiological state of a culture during steady state might differ depending on how it was reached and different properties (e.g. cell and metabolite concentration) can be observed even if the same dilution rate and cultivation conditions are used [38–45]. Such multiplicity of steady states is likely a consequence of toxic waste product accumulation. Lower waste product secretion and higher cell densities indicate a metabolic switch to an energetically more efficient metabolism. This phenomenon could explain the different cell densities and exchange rates observed for three dilution rates (Fig S7 and Fig 5). However, not enough data was available to investigate this phenomenon in more detail.

In literature there is only a small amount of data for mammalian maintenance energy and no data for CHO. Mouse cells require a maintenance energy of 1.7 × 10^−11^ mmol cell^−1^ d^−1^ (65% of the total energy produced at the highest growth rate) [34], which corresponds to 1.1 mmol g^−1^ h^−1^ with a mouse cell dry mass (660 pg/cell) or 2.4-3.6 with a range of CHO dry masses [13]. However, the analysis in [34] was quite simplified. Even though they cultivated the cells with a hydrolysate, they only considered glucose as the energy source and calculated the generated ATP from the secretion rates of lactate and CO_2_. However, mammalian cells in culture will also use glutamine and other AAs as energy source [46].

Depending on the experimental/computational methods used, maintenance energy of cancer cells was estimated to be within 1.6 and 3.7mmolg^−1^ h^−1^ [47,48]. The values were converted from the original publication with CHO specific volume and dry mass values [13].

In other organisms, estimated/measured values widely vary and often depend on the cultivation conditions. In bacteria, the reported values range between 3.15-18.5 [30,49–51], in yeast between 0.44-2.81 mmol g^−1^ h^−1^ [32,52–54]. However, not only does mATP change across organisms and conditions but also during the batch as it is influenced by stress responses [31].

Few other studies also tried to improve the predictions by iCHO1766. For example, Lularevic et al. [55] reduced variability in flux variability analysis by adding carbon availability constraints. In another study, the predictions of intracellular fluxes were improved by adding constraints based on enzyme kinetic information [9]. This also lead to a correct prediction of the overflow metabolism (the secretion of lactate). Together these studies, including the current one, show that adding more constraints to the models is necessary to fully capture cellular metabolism and leads to better predictions. Further developments and a combination of different approaches might lead to further improvement.

## Conclusion

In this work we evaluated the prediction accuracy of CHO GSMM with pFBA. The intracellular fluxes were largely underestimated due to low energy demand and the missing non-growth associated maintenance energy was identified as the main reason for the bad flux predictions. The computationally estimated maintenance energy largely improved the predictions of central carbon metabolism and it was consistent with experimentally determined maintenance energy and with literature values for other mammalian cell lines. Adding this simple constraint to the model leads to a big improvement in the flux prediction accuracy and should not be neglected in constraint-based metabolic modeling of CHO.

## Acknowledgement

This work has been supported by the PhD program BioToP of the Austrian Science Fund (FWF Project W1224) and the COMET center: acib: Next Generation Bioproduction is funded by BMK, BMDW, SFG, Standortagentur Tirol, Government of Lower Austria und Vienna Business Agency in the framework of COMET - Competence Centers for Excellent Technologies. The COMET-Funding Program is managed by the Austrian Research Promotion Agency FFG.

## Supporting information

**Table S1.**
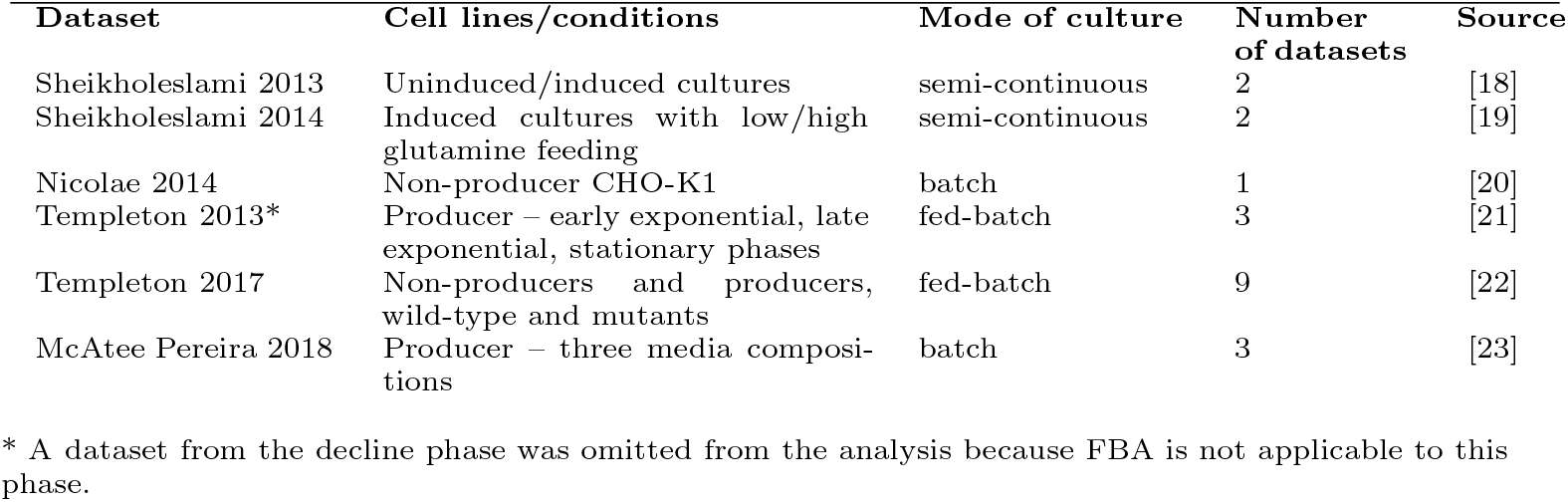
Overview of the analysed ^13^C MFA datasets.

**Fig S1.**
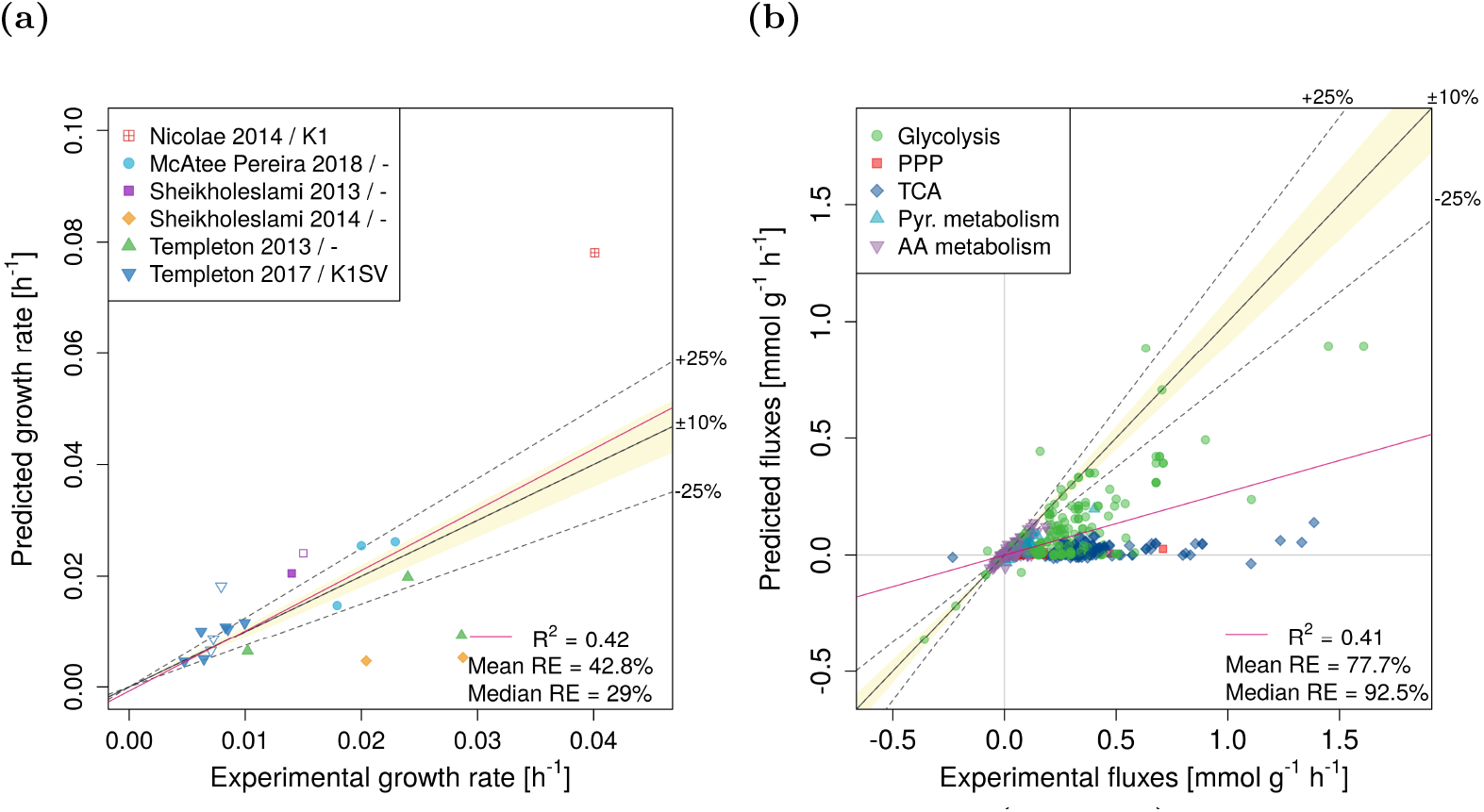
Experimental vs. predicted growth rates (panel a) and intracellular fluxes (panel b). Data is shown for biomass equation `R_biomass_cho_producing` as the objective function. RE – relative error. The legend in panel a indicates the publication and the used CHO cell line (if the information was available). Empty symbols indicate non-producers

**Fig S2.**
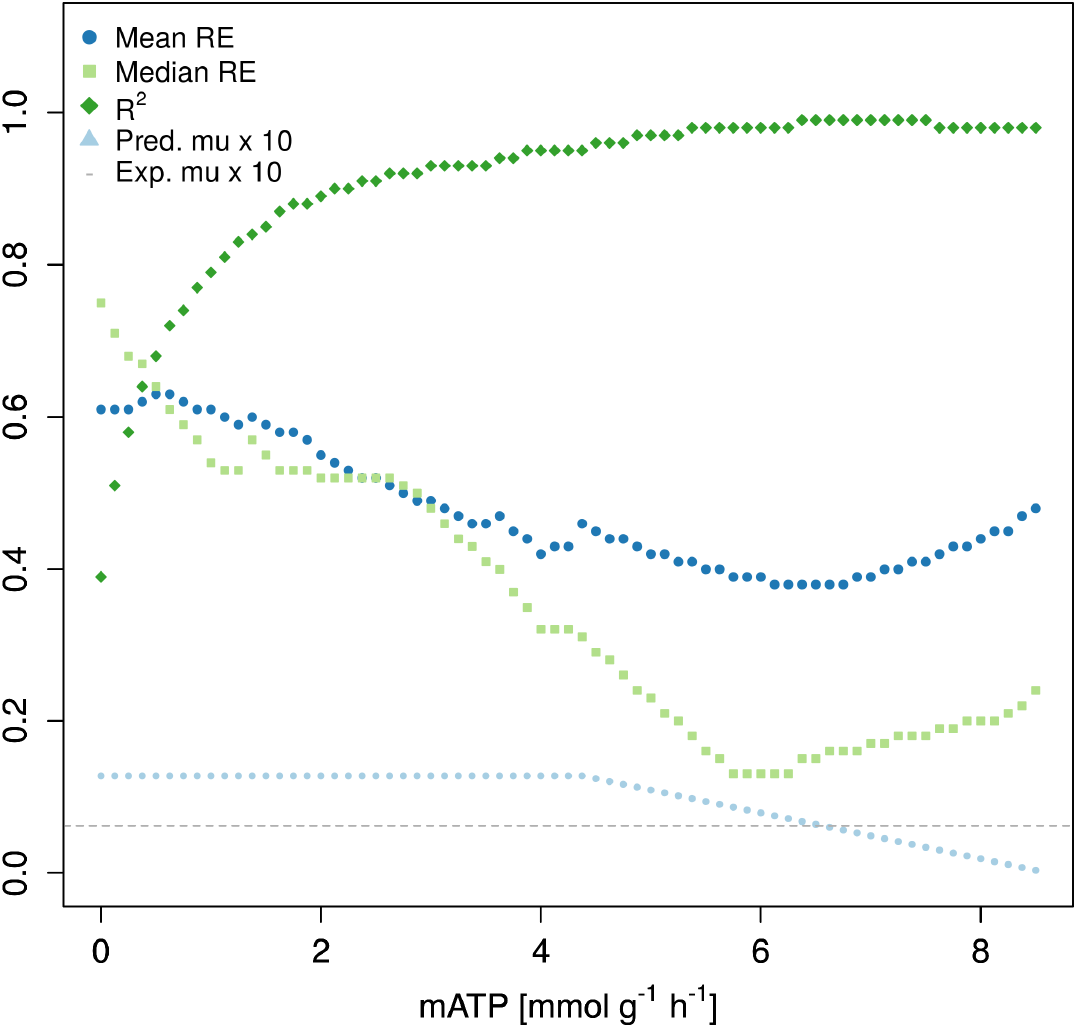
An example of the computational estimation of mATP. mATP was gradually increased and the agreement between experimental and predicted fluxes was evaluated at each step. The mATP value that lead to the smallest median relative error of the fluxes was chosen as the optimal value. Data is shown for the dataset SV-M3 from Templeton 2017 [22] for biomass equation `R_biomass_cho`.

**Fig S3.**
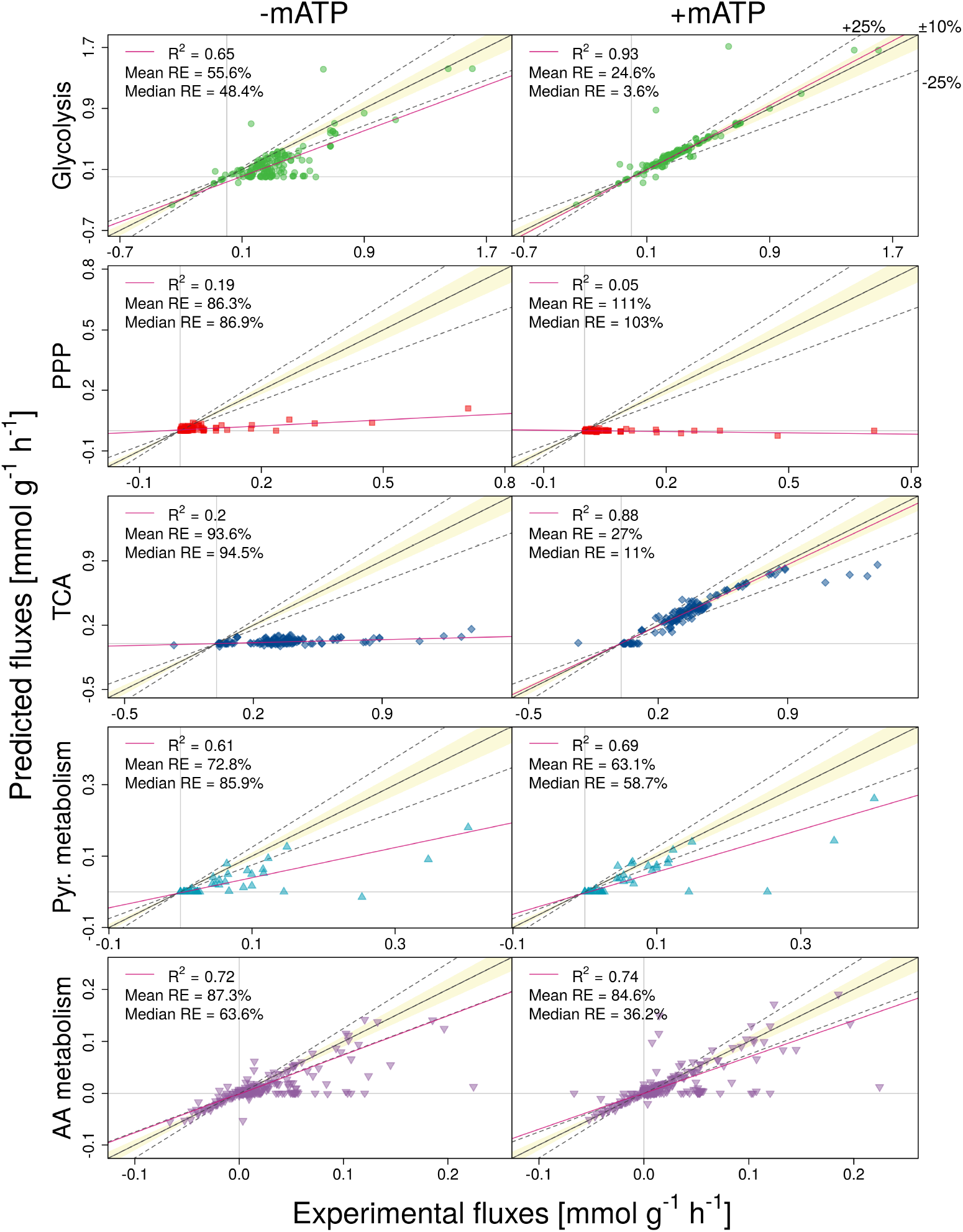
Predictions of intracellular fluxes for the individual subsystems without (-mATP) or with mATP (+mATP) as constraint. Results are shown for `R_biomass_cho` as the objective function. RE – relative error.

**Fig S4.**
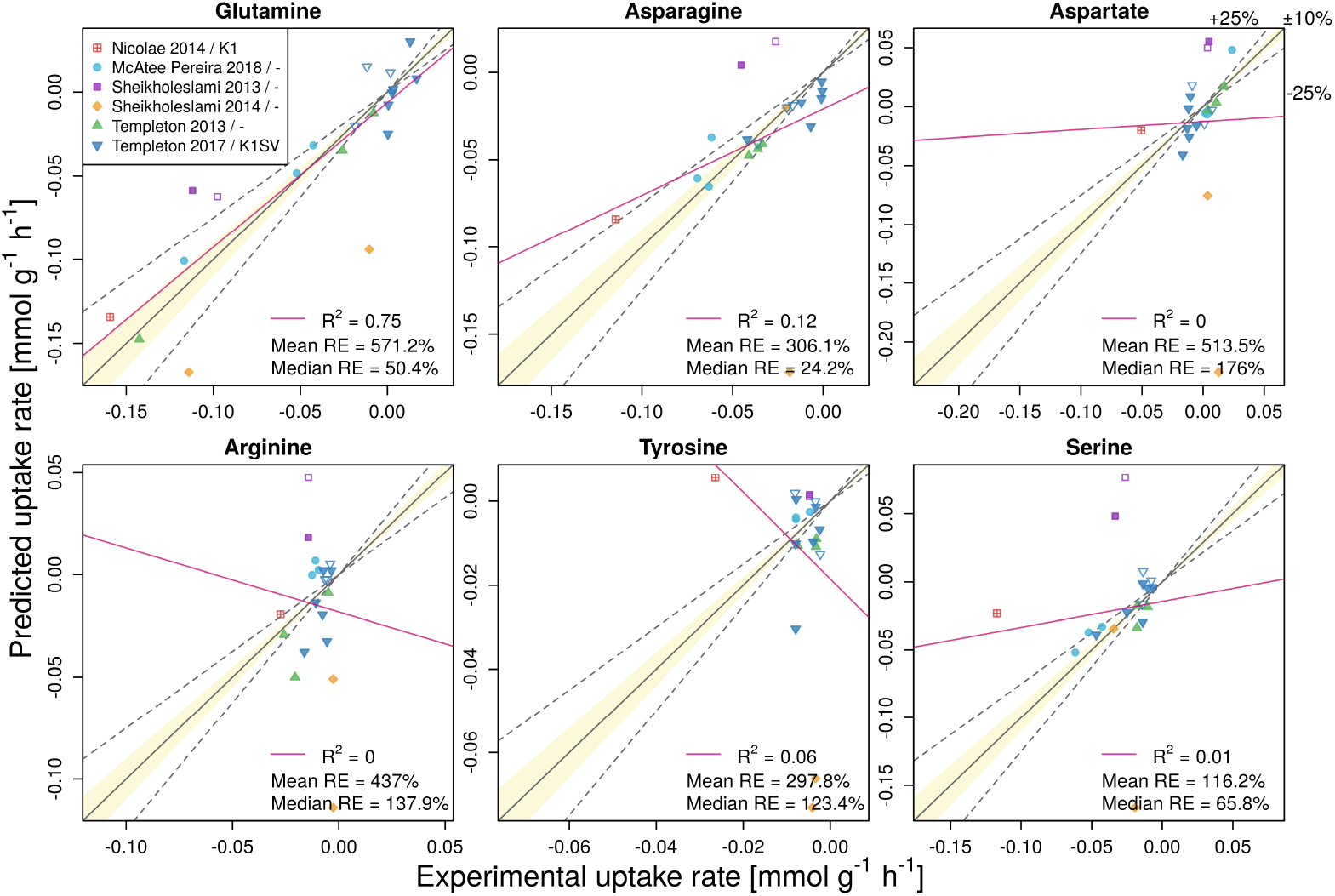
Predictions with different uptake rates as objective functions. Results are shown for R biomass cho as the biomass reaction. RE – relative error. The legend indicates the publication and the used CHO cell line (if the information was available). Empty symbols indicate non-producers

**Fig S5.**
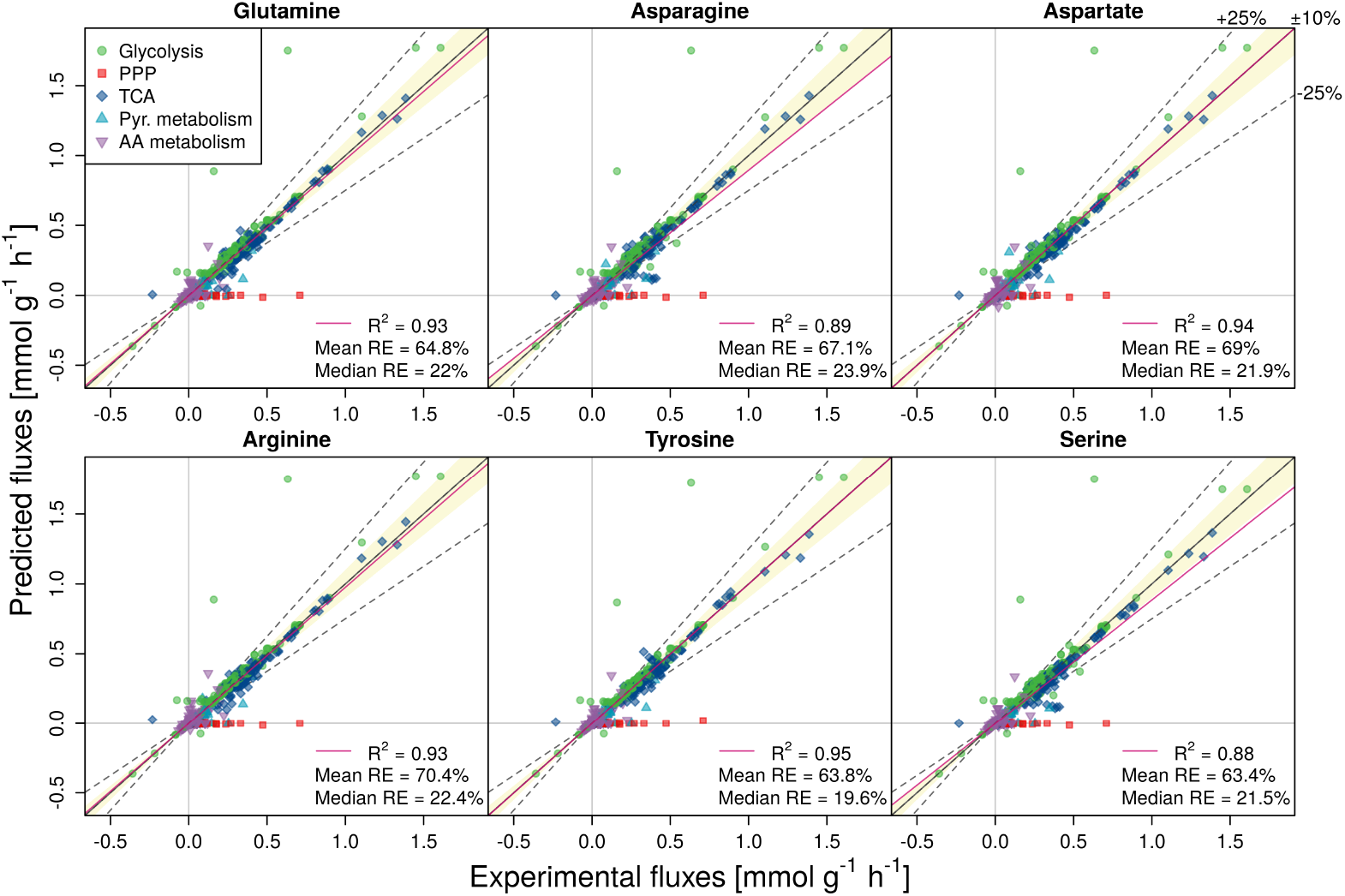
Experimental vs. predicted intracellular fluxes using minimization of nonessential uptakes as objectives. Results are shown for R biomass cho as the biomass reaction. RE – relative error.

**Fig S6.**
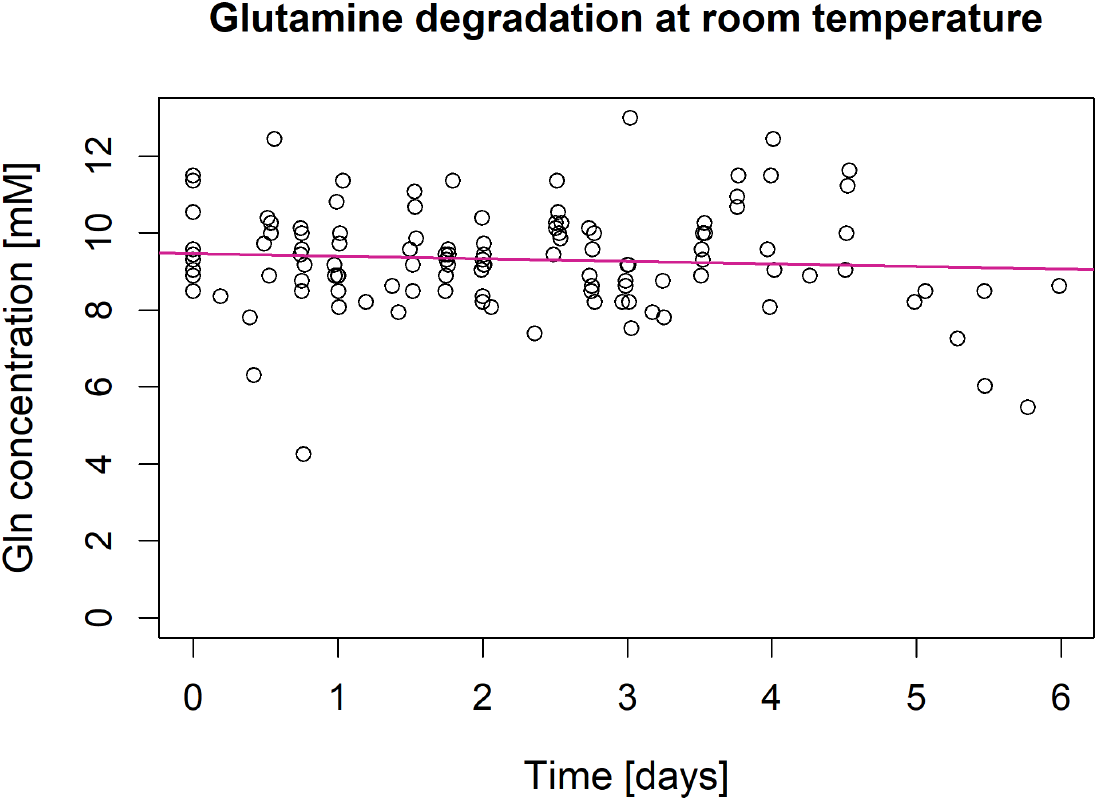
Glutamine degradation at room temperature. The concentration was measured with Bioprofile 100Plus (NOVA Biomedical, MA, USA). The degradation rate (slope of the linear fit) during this time frame is not significant (p-value = 0.402).

**Fig S7.**
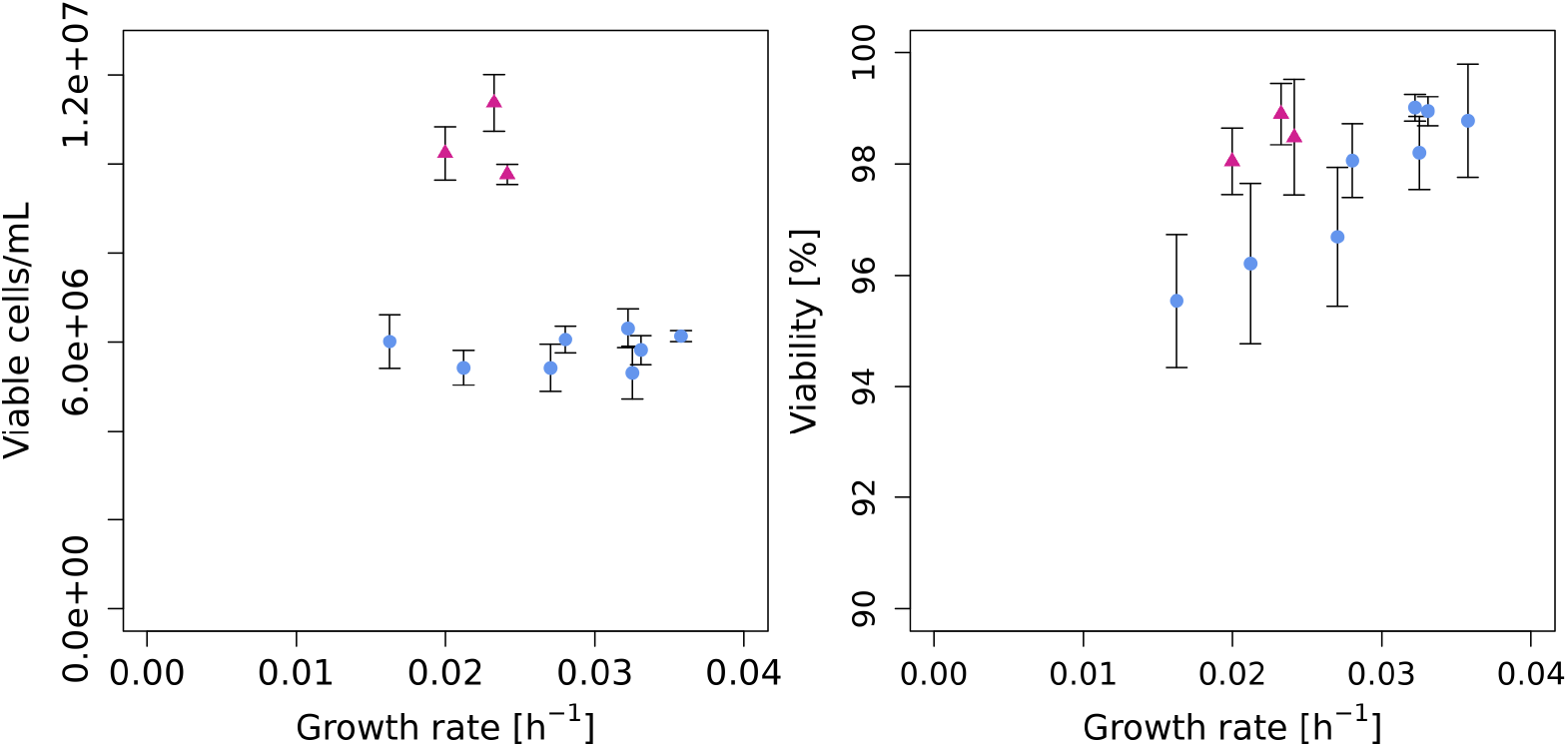
Steady state viable cell density and viability at different growth rates.

