## Supporting information for "Inclusion of maintenance energy improves the intracellular flux predictions of CHO"

**Table S1 . Overview of the analysed $^{13}$C MFA datasets.**

| Dataset | Cell lines/conditions | Mode of culture | Number of datasets | Source |
|---|---|---|---|---|
| Sheikholeslami 2013 | Uninduced/induced cultures | semi-continuous | 2 | [18] |
| Sheikholeslami 2014 | Induced cultures with low/high glutamine feeding | semi-continuous | 2 | [19] |
| Nicolae 2014 | Non-producer CHO-K1 | batch | 1 | [20] |
| Templeton 2013* | Producer – early exponential, late exponential, stationary phases | fed-batch | 3 | [21] |
| Templeton 2017 | Non-producers and producers, wild-type and mutants | fed-batch | 9 | [22] |
| McAtee Pereira 2018 | Producer – three media compositions | batch | 3 | [23] |

* A dataset from the decline phase was omitted from the analysis because FBA is not applicable to this phase.

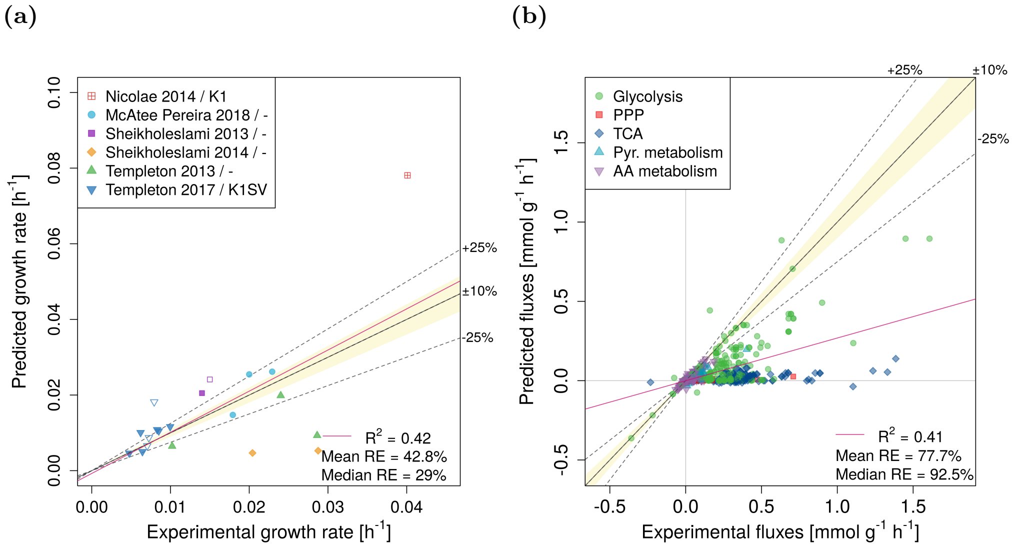


**Fig S1 . Experimental vs. predicted growth rates (panel a) and intracellular fluxes (panel b).** Data is shown for biomass equation `R_biomass_cho_producing` as the objective function. RE – relative error. The legend in panel a indicates the publication and the used CHO cell line (if the information was available). Empty symbols indicate non-producers

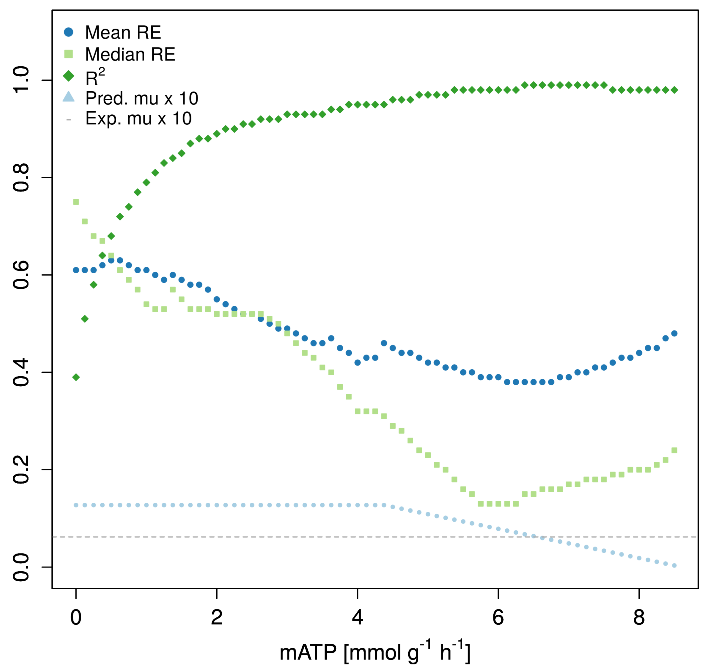


**Fig S2 . An example of the computational estimation of mATP.** mATP was gradually increased and the agreement between experimental and predicted fluxes was evaluated at each step. The mATP value that lead to the smallest median relative error of the fluxes was chosen as the optimal value. Data is shown for the dataset SV-M3 from Templeton 2017 [22] for biomass equation `R_biomass_cho`.

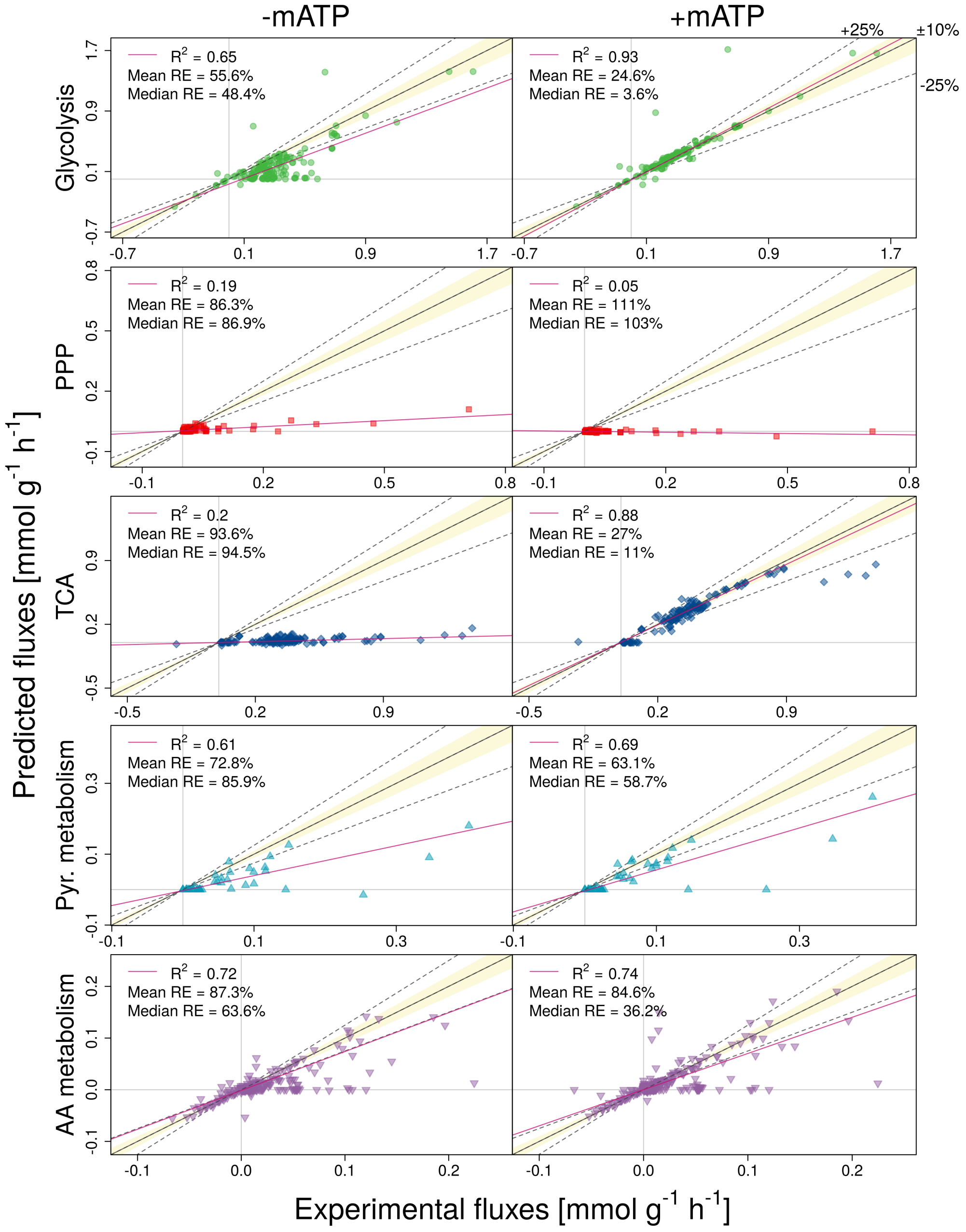


**Fig S3 . Predictions of intracellular fluxes for the individual subsystems without (-mATP) or with mATP (+mATP) as constraint.** Results are shown for `R_biomass_cho` as the objective function. RE – relative error.

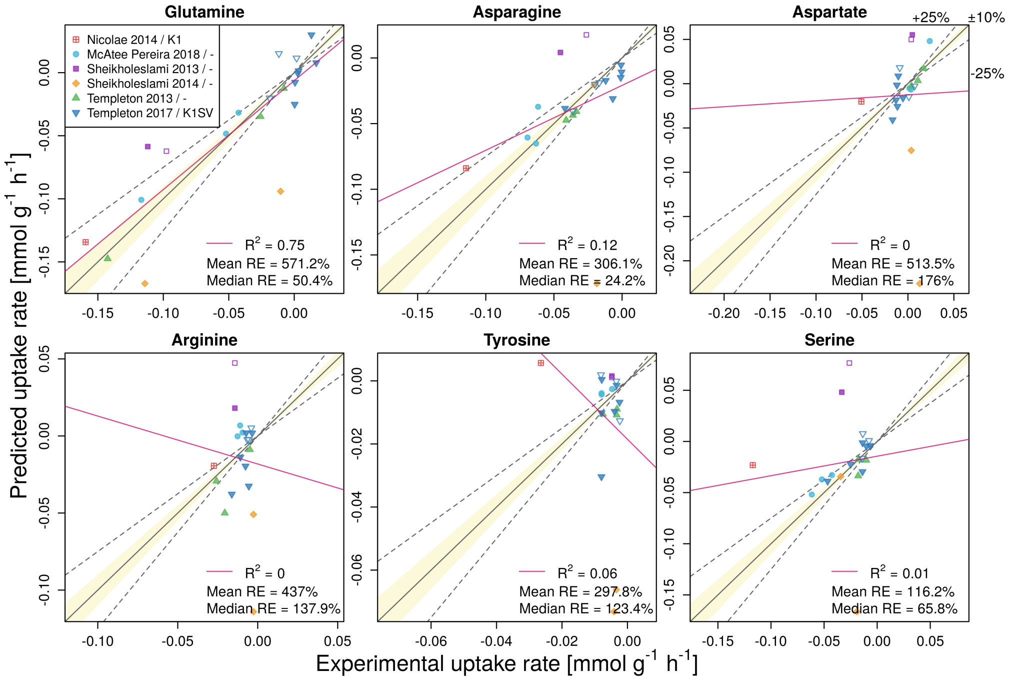


**Fig S4 . Predictions with different uptake rates as objective functions.** Results are shown for `R_biomass_cho` as the biomass reaction. RE – relative error. The legend indicates the publication and the used CHO cell line (if the information was available). Empty symbols indicate non-producers

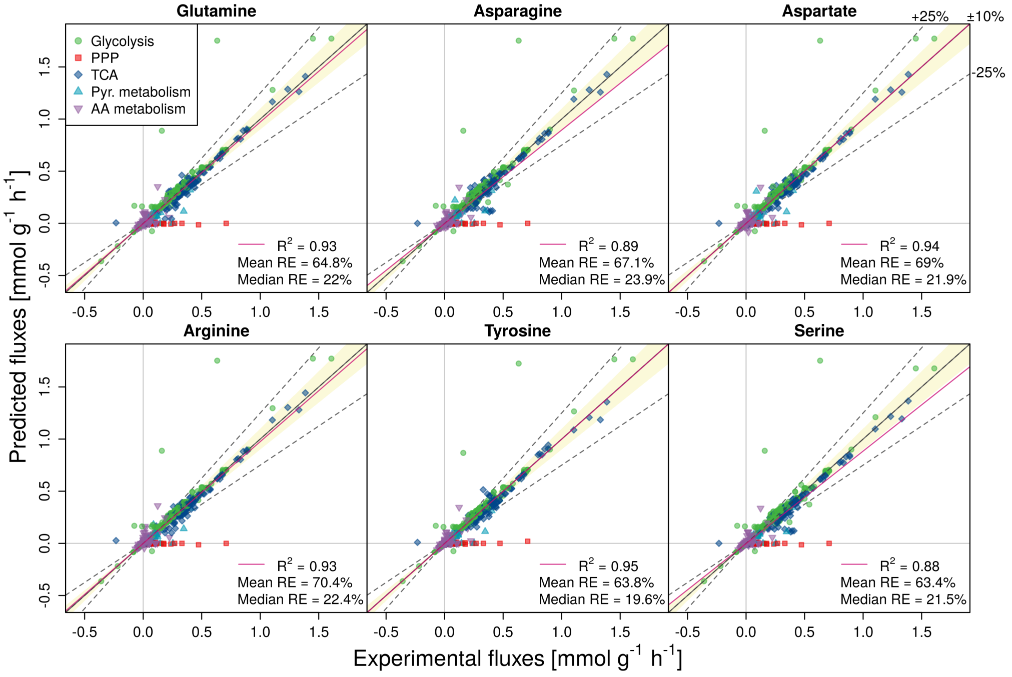


**Fig S5 . Experimental vs. predicted intracellular fluxes using minimization of nonessential uptakes as objectives.** Results are shown for `R_biomass_cho` as the biomass reaction. RE – relative error.

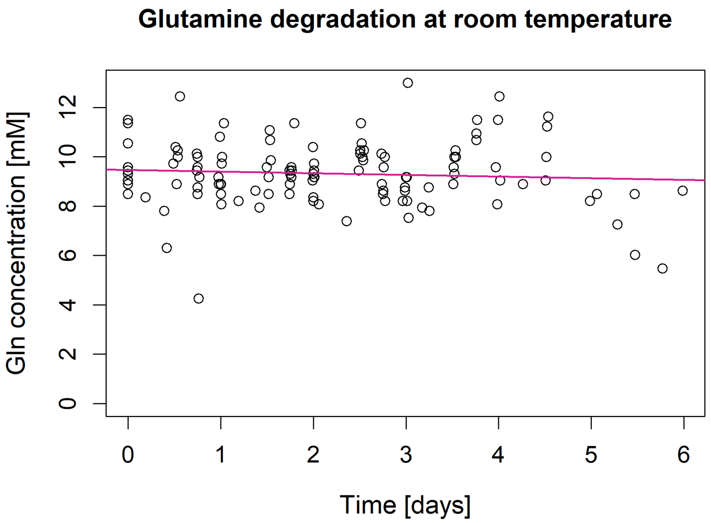


**Fig S6 . Glutamine degradation at room temperature.** The concentration was measured with Bioprofile 100Plus (NOVA Biomedical, MA, USA). The degradation rate (slope of the linear fit) during this time frame is not significant (p-value = 0.402).

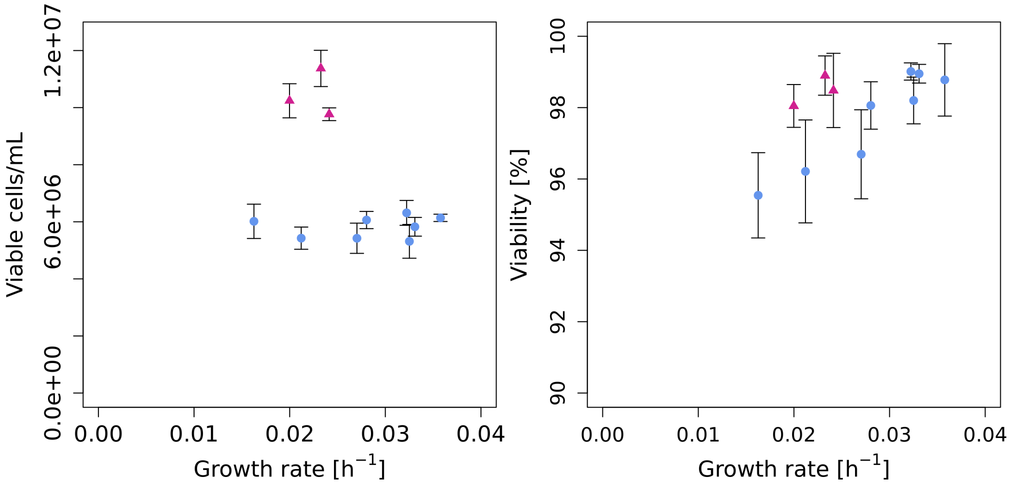


**Fig S7 . Steady state viable cell density and viability at different growth rates.**
